# Larval growth and survival in Indian Butter Catfish, *Ompok bimaculatus* (Bloch): effect of light intensity and photoperiod

**DOI:** 10.1101/623462

**Authors:** Kalpana Arambam, Pradyut Biswas, Soibam Khogen Singh, A. B. Patel, Alok Kumar Jena, Rajkumar Debarjeet Singh, P. K. Pandey

## Abstract

Two sequential indoor rearing trials each of 21 days duration were conducted to investigate the effect of light intensity and photoperiod respectively on the growth and survival of *Ompok bimaculatus* larvae. In first trial, five different light intensities *viz*. 0, 300, 500, 900, 1200 lx were applied randomly to 800 larvae (0.003 g; 0.51 cm) stocked in triplicate following a completely randomized design into aquarium (30.0 x 15.0 x 15.0 cm) tanks. Sequentially, in second trial, five photoperiod cycles (light: dark, L: D) namely, 24L: 0D, 16L: 8D, 12L: 12D, 8L: 16D and 0L: 24D in combination with the best performing light intensity (300 lx) as observed from the first trial were employed in triplicates in similar set up. From the first trial, significantly higher survival was observed in 0 and 300 lx, whereas growth was highest in 900 lx (P < 0.05). In the second trial, survival was higher in continuous darkness (0L: 24D), whereas, maximum growth was recorded in 24L: 0D and 16L: 8D groups (P < 0.05). Performance index (PI) showed no significant difference (P > 0.05) among 0 and 300 lx light intensities, but were reduced at higher light intensities. The lowest PI was found in 12L: 12D and 8L: 16D condition but did not have any effect in other photoperiod cycles. Overall, from the present study it can be concluded that growth of the larvae is found to be higher in higher light intensity (900lx) and longer photoperiodic cycles (24L: 0D and 16L: 8D), however, better survival was recorded in total dark conditions suggesting that continuous dark condition is recommended for better hatchery performance of the larvae.

## Introduction

Indian Butter catfish (*Ompok bimaculatus*), a fish species under Siluridae family has a greater preference among populations in the north-eastern and eastern states of India. The species has good medicinal value, apart from its delicacy in these regions and therefore, considered a priced food fish species fetching market price of US$ 10-12 per kg. The species has considerable reasons for its mass scale promotion in aquaculture. Firstly, the present focus on diversifying available species with those having superior market, particularly the indigenous fishes can benefit upon lucrative return of investment (Pradhan *et al*., 2014) from its culture practices. Secondly, due to the subsequent decline of population in wild, the species has been listed under threatened category (CAMP, 2008). In the midst, successful induced spawning of the said fish is documented earlier (Bhowmick *et al*., 2000; Rawat *et al*., 2018), however, high mortality due to cannibalism is one of the bottlenecks towards successful rearing of the larvae (Chakrabarti *et al*., 2012). Invariably, there is an urgent need for optimise the larval rearing techniques to overcome production of quality seed through manipulated culture conditions.

Among the environmental factors, photoperiod is considered the most important bio-factor influencing the growth and survival of the fishes (Head and Malison, 2000; Downing and Litvak, 2001; Ruchin, 2004; Downing, 2002; Giri *et al*., 2002). For instance, photoperiods longer than that of ambient conditions increased growth of larval rabbit fish, *Siganus guttatus* (Duray and Kohno, 1988), sea bass, *Dicentrarchus labrax* (Barahona-Fernandes, 1979; Ronzani Cerqueira *et al*., 1991), barramundi, *Lates calcarifer* (Barlow *et al*., 1995), greenback flounder, *Rhombosolea tapirina* (Hart *et al*., 1996), freshwater sheat-fish, *Wallago attu* (Giri *et al*., 2002) and european sea bass, *Dicentrarchus labrax* (Villamizar *et al*., 2009). Besides, shorter day lengths are even preferable for larvae [e.g., striped bass *Morone saxatilis* (Martin-Robichaud and Peterson, 1998). Although light helps to be the primary sense involved in foraging activity and feeding (Puvanendran and Brown, 2002), but also plays a key roles in internal synchronization for the rhythmic synthesis and release of time-keeping hormones (i.e. melatonin), whose signal affects rhythmic physiological functions in fish (Bromage *et al*., 2001; Amano *et al*., 2003; Falcon *et al*., 2010; Migaud *et al*., 2010). However, the optimal photoperiod for larval growth and survival may differ, and also vary with larval ontogeny (Fielder *et al*., 2002).

Further, in aquaculture, finding the correct environmental lighting for most of teleost larvae is complex. For example, it is established that most fish larvae need a minimal threshold of light intensity to survive. This threshold light intensity and periodicity acts a major role in regulating fish larval activity (e.g. feeding), prey selection and localization by fish (Nwosu and Holzlohner, 2000). Light intensity can also influence the fish to stress (Strand *et al*., 2007; Rotllant *et al*., 2003; Papoutsoglou *et al*., 2005), which may affect their behaviour by altering swimming performance, activity levels and habitat consumption (Mesa and Schreck, 1989; Schreck *et al*., 1997). For example, African catfish *(Clarias gariepinus)* juveniles showed the higher activity in 150 lux, compared to 15 lux, was considered a stress reaction (Almazán-Rueda *et al*., 2004). The lighting regime has been documented in many cultured species by fish farmers to get benefits in terms of survival and growth under controlled rearing conditions (Tandler and Helps, 1985; Batty, 1987; Downing and Litvak, 1999). In contrast, standard lighting systems usually used in hatcheries create bright point light sources that are neither environment-specific nor species-specific and can potentially compromise fish welfare (Villamizar *et al*., 2009). Moreover, light intensity and photoperiod can both control the ability for larvae to inflate their swim bladders in terms associated with larval survival (Battaglene, 1995; Fielder *et al*., 2002).

Considering the lack of research available regarding the application of optimal light intensity and photoperiod manipulation in *O. bimaculatus* larviculture, the present work was taken up to delineate its effect on growth performance and survival of the species under controlled environmental conditions. The present study will be helpful in establishing an optimal environmental photo-periodic and light intensity regime for *O. bimaculatus* larviculture as already established in other catfishes of commercial importance.

## Results

### Water quality

The mean values of the physiochemical parameters of water in both the experiments show variations among the treatments, however; not significantly different (P > 0.05). Mean temperature ranged from 25.90 to 28.80 °C; dissolved oxygen levels was above 6 mg L^−1^; pH ranged between 7.2 to 7.8; total alkalinity ranged between 53.9-61.2 mg CaCO_3_ L^−1^; total hardness ranged between 35.1–42.6 mg CaCO_3_ L^−1^; ammonia nitrogen (NH_3_^+^-N), nitrate nitrogen (NO_3_^−^-N) and nitrite nitrogen (NO_2_^−^-N) ranged between 0.01–0.12 mg L^−1^, 0.05–0.21 mg L^−1^ and 0.01–0.05 mg L^−1^ respectively. The above recorded water quality parameter values during the experimental groups fell within the standard optimal range for fish culture (Boyd, 1982).

### Trial 1: light intensities effect on growth and survival

The growth performance of the larvae were significantly affected (P < 0.05) by the experimental light intensities provided (**Table 1**). The final weight (g), weight gain (%) and specific growth rate (SGR; % day^−1^) was found significantly higher (P < 0.05) in 900 lx group compared to others, whereas, significantly lower value was found in 0 lx light intensity groups (P<0.05). Final length and length increment (cm) was significantly lower in 0 lx groups while, no significant changes was observed among other groups. Other yield parameters such as survival (%), total biomass (g) and performance index (PI) during the length of the study were depicted in **Table 2**. The highest (P < 0.05) survival and performance index were found in 0 lx and 300 lx light intensities, however, no significant difference was observed among 500lx, 900lx and 1200 lx treatments. Significantly higher biomass (g) was recorded in lower intensities treatments (i.e. 0 lx to 500 lx) and relatively lower value was found in the higher light intensities (P<0.05).

**Table 1.** Growth parameters of *Ompok bimaculatus* larvae reared under different light intensities

| Parameter | Light intensities (lx) |  |  |  |  | P-values |  |  |
| --- | --- | --- | --- | --- | --- | --- | --- | --- |
|  | 0lx | 300lx | 500lx | 900lx | 1200lx | Overall | Linear | Quadratic |
| **Initial length(cm)** | 0.52±0.05 | 0.49±0.03 | 0.53±0.01 | 0.53±0.04 | 0.54±0.01 | 0.832 | 0.388 | 0.757 |
| **Initial weight(g)** | 0.0032±0.0001 | 0.0032±0.00002 | 0.0031±0.00003 | 0.0031±0.00002 | 0.0032±0.00008 | 0.357 | 0.405 | 0.073 |
| **Final length(cm)** | 2.27±0.04<sup>a</sup> | 2.50±0.05<sup>b</sup> | 2.53±0.02<sup>b</sup> | 2.52±0.05<sup>b</sup> | 2.53±0.05<sup>b</sup> | 0.006 | 0.002 | 0.012 |
| **Final weight(g)** | 0.067±0.002<sup>a</sup> | 0.08±0.003<sup>b</sup> | 0.08±0.003<sup>b</sup> | 0.09±0.004<sup>c</sup> | 0.08±0.002<sup>b</sup> | 0.001 | 0.000 | 0.006 |
| **Body weight gain (%)** | 1955.76±28.15<sup>a</sup> | 2356.58 ±101.34<sup>b</sup> | 2540.83<br>±107.05<sup>b</sup> | 2856.18±118.86<sup>c</sup> | 2476.10<br>±72.26<sup>b</sup> | 0.001 | 0.000 | 0.002 |
| **Length increment (cm)** | 1.75±0.04<sup>a</sup> | 2.01±0.04<sup>b</sup> | 2.00 ±0.01<sup>b</sup> | 1.99±0.05<sup>b</sup> | 1.99±0.04<sup>b</sup> | 0.003 | 0.003 | 0.004 |
| **Specific growth rate (%**<br>**d<sup>-1</sup>)** | 3.02±0.01<sup>a</sup> | 3.20 ±0.04<sup>b</sup> | 3.27 ±0.04<sup>b</sup> | 3.38±0.05<sup>c</sup> | 3.25 ±0.03<sup>b</sup> | 0.000 | 0.000 | 0.001 |
| **Mean daily weight gain**<br>**(%)** | 0.42±0.01<sup>a</sup> | 0.41±0.02<sup>a</sup> | 0.44±0.01<sup>a</sup> | 0.50±0.02<sup>b</sup> | 0.44±0.01<sup>a</sup> | 0.030 | 0.045 | 0.313 |

\*Overall mean value (± SE, n = 3) having different superscript in the same row shows significance difference (P < 0.05); a, b and c denotes significance differences between different treatments.

**Table 2.**
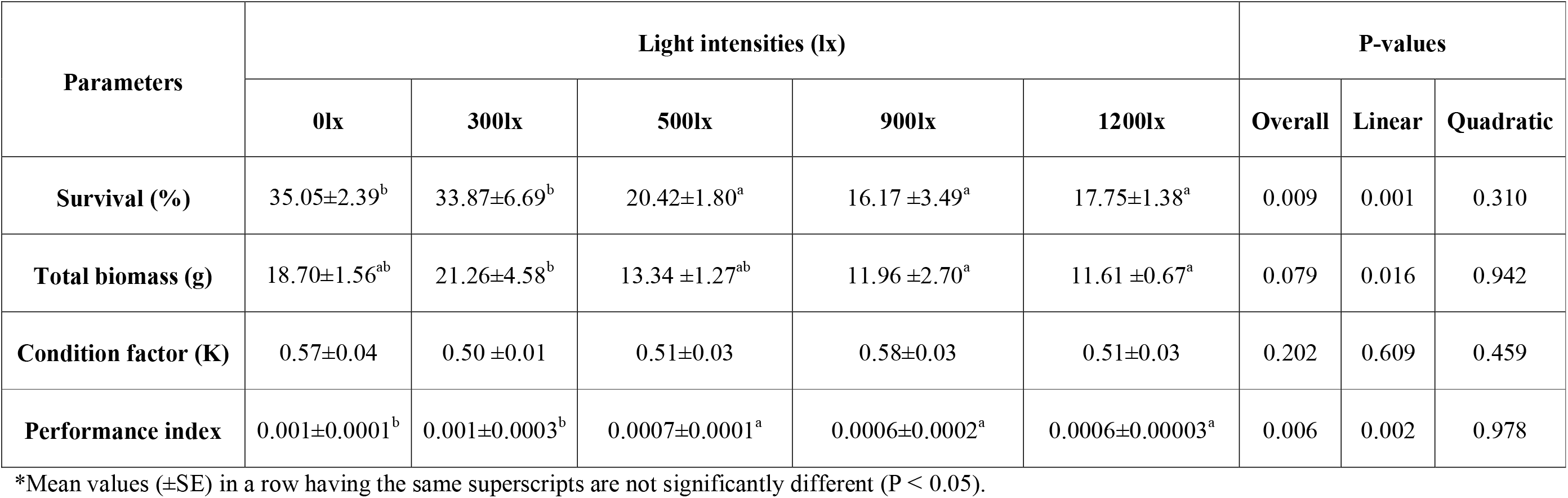
Yield parameters of *Ompok bimaculatus* larvae reared under different light intensities

| Parameters | Light intensities (lx) |  |  |  |  | P-values |  |  |
| --- | --- | --- | --- | --- | --- | --- | --- | --- |
|  | 0lx | 300lx | 500lx | 900lx | 1200lx | Overall | Linear | Quadratic |
| Survival (%) | 35.05±2.39<sup>b</sup> | 33.87±6.69<sup>b</sup> | 20.42±1.80<sup>a</sup> | 16.17 ±3.49<sup>a</sup> | 17.75±1.38<sup>a</sup> | 0.009 | 0.001 | 0.310 |
| Total biomass (g) | 18.70±1.56<sup>ab</sup> | 21.26±4.58<sup>b</sup> | 13.34 ±1.27<sup>ab</sup> | 11.96 ±2.70<sup>a</sup> | 11.61 ±0.67<sup>a</sup> | 0.079 | 0.016 | 0.942 |
| Condition factor (K) | 0.57±0.04 | 0.50 ±0.01 | 0.51±0.03 | 0.58±0.03 | 0.51±0.03 | 0.202 | 0.609 | 0.459 |
| Performance index | 0.001±0.0001<sup>b</sup> | 0.001±0.0003<sup>b</sup> | 0.0007±0.0001<sup>a</sup> | 0.0006±0.0002<sup>a</sup> | 0.0006±0.00003<sup>a</sup> | 0.006 | 0.002 | 0.978 |

\*Mean values (±SE) in a row having the same superscripts are not significantly different (P < 0.05).

### Trial 2: photoperiod effect on growth and survival

The growth and yield parameters of larvae under different treatments are shown in **Table 3 and 4** respectively. The growth performance of larvae was significantly affected (P < 0.05) by varied photoperiod cycles. The initial length and weight of *Ompok bimaculatus* larvae, stocked in all the treatments were the same (P > 0.05). The highest growth (P < 0.05) was observed in the treatment with 24L: 0D and 16L: 8D photoperiods and no significant difference (P > 0.05) was found among 12L: 12D, 8L: 16D and 0L: 24D treatments. The final mean length and length increment of larvae did not differ significantly (P > 0.05) among the groups. Moreover, the highest survival (%) was recorded in 0L: 24D and the lowest in 24L: 0D photoperiod cycles. The total biomass was found higher in 24L: 0D, 16L: 8D and 0L: 24D conditions but did not significantly differ among the groups. Significantly lower condition factor was reported in 0L: 24D photoperiods while, the performance index showed linearly higher in 24L: 0D, 16L: 8D and 0L: 24D photoperiod cycles.

**Table 3.** Growth performance of *Ompok bimaculatus* larvae reared under different photoperiods.

| Parameters | Photoperiods |  |  |  |  | P-values |  |  |
| --- | --- | --- | --- | --- | --- | --- | --- | --- |
|  | 24L:0D | 16L:8D | 12L:12D | 8L:16D | 0L:24D | Overall | Linear | Quadratic |
| **Initial length(cm)** | 0.4833±0.02 | 0.5033± 0.03 | 0.5133±0.03 | 0.5400±0.01 | 0.5160±0.05 | 0.731 | 0.290 | 0.524 |
| **Initial weight(g)** | 0.003 ± 0.0001 | 0.003 ±0.00005 | 0.003 ± 0.00007 | 0.003 ±0.00007 | 0.003 ± 0.00006 | 0.283 | 0.067 | 0.334 |
| **Final length(cm)** | 2.05 ± 0.07 | 2.27 ± 0.04 | 2.01 ± 0.21 | 1.96 ± 0.04 | 2.27 ± 0.03 | 0.177 | 0.712 | 0.374 |
| **Final weight(g)** | 0.091± 0.01<sup>b</sup> | 0.086± 0.01<sup>b</sup> | 0.061 ± 0.004<sup>a</sup> | 0.063±0.004<sup>a</sup> | 0.064± 0.001<sup>a</sup> | 0.005 | 0.001 | 0.078 |
| **Body weight gain (%)** | 2907.76± 179.93<sup>b</sup> | 2722.66 ± 251.71<sup>b</sup> | 1901.84± 146.07<sup>a</sup> | 1964.15± 193.48<sup>a</sup> | 1873.88± 56.03<sup>a</sup> | 0.004 | 0.001 | 0.137 |
| **Length increment (cm)** | 1.563 ±0.09 | 1.77 ± 0.05 | 1.50 ± 0.20 | 1.42 ± 0.03 | 1.751 ± 0.04 | 0.133 | 0.941 | 0.279 |
| **Specific growth rate (% d<sup>-1</sup>)** | 3.40±0.06<sup>b</sup> | 3.33±0.09<sup>b</sup> | 2.99± 0.07<sup>a</sup> | 3.02± 0.10<sup>a</sup> | 2.98± 0.03<sup>a</sup> | 0.004 | 0.001 | 0.150 |
| **Mean daily weight gain (%)** | 0.58± 0.05<sup>b</sup> | 0.55± 0.04<sup>b</sup> | 0.39± 0.03<sup>a</sup> | 0.40±0.03<sup>a</sup> | 0.40± 0.01<sup>a</sup> | 0.005 | 0.001 | 0.080 |

\*Mean values (± SE, n = 150; r=3) in a row having the same superscripts are not significantly different (p ≥0.05).

**Table 4.** Yield parameters of *Ompok bimaculatus* larvae reared under different photoperiods

| Parameters | Photoperiods |  |  |  |  | P values |  |  |
| --- | --- | --- | --- | --- | --- | --- | --- | --- |
|  | 24L:0D | 16L:8D | 12L:12D | 8L:16D | 0L:24D | Overall | Linear | Quadratic |
| Survival (%) | 23.96 ± 2.29<sup>a</sup> | 28.63±2.49<sup>ab</sup> | 28.21±1.80<sup>ab</sup> | 27.96±1.55<sup>ab</sup> | 35.05 ± 2.39<sup>c</sup> | 0.050 | 0.010 | 0.545 |
| Total biomass (g) | 17.42± 2.38<sup>ab</sup> | 19.41 ± 0.31<sup>b</sup> | 13.67 ± 0.83<sup>a</sup> | 14.24 ± 1.76<sup>a</sup> | 17.94 ± 1.52<sup>ab</sup> | 0.102 | 0.417 | 0.123 |
| Condition factor (K) | 1.06±0.09<sup>b</sup> | 0.73±0.09<sup>ab</sup> | 0.81±0.19<sup>ab</sup> | 0.85±0.11<sup>ab</sup> | 0.55±0.03<sup>a</sup> | 0.097 | 0.030 | 0.981 |
| Performance index | 0.0010±0.0001<sup>ab</sup> | 0.0011±0.00001<sup>b</sup> | 0.0007±0.00004<sup>a</sup> | 0.0008±0.0001<sup>a</sup> | 0.0010±0.00008<sup>ab</sup> | 0.095 | 0.038 | 0.121 |

\*Different superscripts in the same row signify statistical differences (p<0.05) (mean ± S.E.).

## Discussion

The present study showed that the growth performance of Ompok bimaculatus larvae was significantly affected by light intensities. The specific growth rate, body weight gain (%) and mean daily weight gain (%) was found significantly higher in 900 lux light intensities and lower in 0 lux groups whereas, no significance response was reported in other groups (Table 1). The better growth performance of the larvae at medium light intensities (i.e. 900 lux) was attributed by increased feed intake due the excitation of retinal pigments may permit for the contrast between prey and background (Downing and Litvak, 2001; Strand et al., 2007; Villamizar et al., 2011). Moreover, the higher light intensity may also influences the behavioural changes of both larvae and prey resulting in increased encounter rates (Downing and Litvak, 1999). Several other studies also reported that a threshold light intensities increased the growth in fishes such as spotted sand bass, *Paralabrax maculatofasciatus* (Downing and Litvak, 2001; Brown *et al*., 2003; Pena *et al*., 2004; Villamizar *et al*., 2011) which is relevant to the present study. However, in the present study the growth performance of the *Ompok bimaculatus* larvae had declined in elevated light intensity (i.e. 1200 lx). This finding may be influenced by the environmental stress at the elevated light intensities resulting in lower feeding rates (Delabbio *et al*., 2015). Similar findings were also reported in the Atlantic cod larvae *Gadus morhua* (Puvanendran and Brown, 1998) and European sea bass (Barahona-Fernandes, 1979) where light intensities beyond the threshold limit had a negative effect on the growth performance of the fish. The complete dark groups (0 lx) showed significantly lower final length and the length increment while, no significant response (P>0.05) was observed in others. This result was possibly due to lower growth rate in 0 lx groups (**Table 1**).

This study also revealed that fish reared under 0 lx (35.05±2.39%) and 300 lx (33.87±6.69%) light intensities showed significantly highest (P<0.05) survival (%), whereas no significant difference (P > 0.05) was recorded in other groups (**Table 2**). Similar observation was reported in Atlantic cod larvae *Gadus morhua* (Puvanendran and Brown, 2002), Atlantic halibut *Hippoglossus hippoglossus* (Bolla and Holmefjord, 1988), African catfish *Clarias geripineus* (Britz and Pienaar, 1992) and Orange-Spotted Grouper *Epinephelus coioides* (Takeshita and Soyano, 2008) where low light intensities (0 and 300 lx) had lower mortality rates when compared to larvae reared at higher light intensities. This result may be due to the fact that an elevation of light intensity acts as environmental stressor that induces swim bladder hypertrophy which possibly leads to high mortality (Johnson and Katavic, 1984; Strand *et al*., 2007; Villamizar *et al*., 2011). Moreover, the light intensity known to persuade the secretion of retina and pineal gland (light receptor) in fish (Bromage *et al*., 2001; Amano *et al*., 2003; Falcon *et al*., 2010). As a result melatonin release increased with the dark and decreased with the light conditions in the rainbow trout and goldfish (Iigo and Aida, 1995) which reported to inhibit the cannibalism of *Aequidens pulcher* (Munro, 1986). This finding may be possibly the reason behind the decreased cannibalism of Pabda larvae reared under lower light intensities (i.e. 0 and 300 lx). The present study was inconformity with the observation in African catfish *Clarias geripineus* (Britz and Pienaar, 1992), yellowtail (Sakakura and Tsukamoto, 1997) and Orange-Spotted Grouper *Epinephelus coioides* (Takeshita and Soyano, 2008) where higher light intensity showed increasing incidence of territorial aggression leads to mortality. The result of the study also showed that total biomass and performance index was significantly higher (P < 0.05) in lower light intensity groups followed by other groups, whereas, lower value was recorded in higher intensity groups (i.e. 900 and 1200 lx). This may be due to more survivability in case of lower light intensity groups which indicate the optimum light intensity for the *O. bimaculatus* larvae.

It has been reported that freshwater fish species are more responsive to photoperiod than marine and diadromous species (El-Sayed *et al*., 2004). Moreover, the effect of marine species to photoperiods has been well documented, while little information is available on freshwater species. The results of the present study indicated that the *Ompok bimaculatus* larvae were clearly affected by the different rearing conditions of photoperiod displaying better growth (wet weight), specific growth rate, weight gain (%) and daily mean weight gain (%) in long light cycles compared to those exposed to intermediate or short light periods. A similar results was also observed with several fish larvae such as obscure puffer *Takifugu obscurus* larvae (Shi *et al*., 2010), freshwater cat fish *Wallgo attu* (Giri *et al*., 2002), *Clarias gariepinus* (Ataguba *et al*., 2015) and rabbitfish *Siganus guttatus* (Duray and Kohno, 1988), where the better growth rate was achieved with increasing light periods. This outcome may be due to the fact that continuous light offered an increased chance for the larvae to come across food organisms (Shi *et al*, 2010), and the darkness reduced the growth, possibly by hindering the intake of an adequate amount of food (Giri *et al*., 2002; Hart *et al*., 1996) which is relevant to the present study. It also reported that in shorter light periods may require more energy to synchronizing an endogenous rhythm to the external environment (Biswas and Takeuchi, 2002; Biswas *et al*., 2002; El-Sayed *et al*., 2004) and more standard metabolic rate (Biswas *et al*., 2002), which leads to decline in somatic growth in fish. Hence, it is concluded that *Ompok bimaculatus* conserve energy when reared under different photoperiod cycles with longer light phases. The survival of *Ompok bimaculatus* larvae was significantly influenced by photoperiods. In the present study, Pabda larvae rearing under continuous dark condition showed maximum survival rates with reduced cannibalism. Similarly, Parameswaran *et al*. (1970) and Chakrabarti (2012) also reported that the dark condition is preferred for better survival of *O. bimaculatus* larvae during in-door rearing as their cannibalistic habit profound in light conditions. The lower survival (%) in continuous light (24L: 0D) photoperiod was possibly due to carnivorous and cannibalistic mode of life of *O. bimaculatus* during early stages and also attributed by the visual clues during the rearing under continuous light (24L: 0D) cycles. This was probably due to the absence of stress and aggressiveness, as well as the suppression of locomotory activities in the dark. The study also showed that total biomass and performance index did not vary with the treatments because of differential growth and survival. Condition factor was significantly higher (P<0.05) in continuous light (24L: 00D) conditions may be due to more chances of cannibalistic encounter among larvae due to light.

In the present study, although, the growth performance of the Pabda larvae was found significantly higher in higher light intensity (i.e. 900lx) and longer light cycles (i.e. 24L: 0D and 16L: 8D), but the performance of a hatchery generally judged based on the survivor rather than the growth. The better survival was recorded in total dark conditions suggesting that the larval rearing in continuous dark condition is recommended for better hatchery performance.

## Materials and methods

### Sources of larvae

The fish attained maturity at the end of the first year. Fully ripe females and males (40g or above) were taken from the pond of College of Fisheries, Lembucherra, Tripura, India and used for induced breeding. Females can be distinguished by a rounder, fuller abdomen, reddish vent colour and rounded genital papilla. Males have an extended and pointed genital papilla. The induce breeding was carried out by administrating a single injection of inducing agent Ovatide^®^ @ 1–1.5 ml kg^−1^ body weight for females and 0.5-1.0ml kg^−1^ body weight for males (Chakrabarti *et al*., 2012). Females were stripped for spawning after 8 – 10 hours of hormone injection and the eggs were collected in a tray. Milt was attained from males by surgically removing the testes, which were macerated to get a suspension to be mixed with the eggs for fertilisation. Then the fertilized eggs were washed carefully with clean water and transferred to a fibreglass tank for hatching, with constant aeration. Hatching takes place within 22-24 hours of fertilization. Subsequently newly hatched larvae (2 days old) were taken for the study.

### Experimental set up

The study was carried out for 21 days at the wet laboratory of College of Fisheries, Central Agricultural University (Imphal), Lembucherra, Tripura, India to find out the effect of light intensity and photoperiod on growth performance, survival and production of *O. bimaculatus* larvae. The larvae were reared in glass aquarium (30.0 x 15.0 x 15.0 cm) filled with 25-L of chlorine-free bore well water with continuous aeration. Aquariums were properly cleaned, sterilised and dried before introducing the larvae. Larvae were fed with chopped fresh tubifex upto satiation thrice everyday at 06:00, 14:00 and 22:00 h. Dead larvae, faeces, and other debris were siphoned out, and the number of dead larvae was recorded every day. Fifty percent of the water in each aquarium was replaced with fresh water daily. The water quality parameters *viz*. temperature, pH, dissolved oxygen (DO), free carbon dioxide (CO_2_), carbonate hardness, ammonia-N and nitrate–N were analysed every weekly interval following the standard procedure (APHA, 2005) in both the experiments. Sampling of fish was done at 5-days intervals and completely randomized design (CRD) was followed throughout the study.

#### Trial 1: Effect of light intensity on growth and survival of larvae

Five light intensity namely 0, 300, 500, 900 and1200 lx were employed in triplicates. Required was obtained by placing incandescent white light (300 lx to 1200 lx; Soft White bulb) 30 cm over head the water surface constantly. Each aquaria were enclosed within a box made from black plastic PVC sheeting to avoid the escape of light to the surrounding tanks. Light intensity was measured at the aquaria water surface using a lux meter (model ZU 104766, AEG, Frankfurt, Germany). Larvae (2-DPH; mean weight 0.003±0.00 g; mean total length 0.52±0.03 cm) were randomly assigned to 15 aquaria ((30.0 x 15.0 x 15.0 cm, 25-L) with 800 nos. of larvae per aquarium filled with chlorine-free bore well water with continuous aeration.

#### Trial 2: Effect of photoperiod on growth and survival of larvae

Larvae were reared in five different photoperiods cycles (light: dark, L: D) at 300lx namely 24 h continuous light (24L: 0D), 16 h light-08 h darkness (16L: 8D), 12 h light-12 h darkness (12L: 12D), 08 h light-16 h darkness (8L: 16D) and 24 h continuous darkness (0L: 24D), using incandescent white bulbs. Light intensity was kept constant at 300 lx throughout the study which was placed at 30 cm vertically from the water surface as described by Daniels et al. (1996). The complete darkness was obtained by covering the aquariums with black plastic PVC sheets with the provision for administrating of feed to the larvae. Photoperiods were maintained by using a 24-h timer (Multi 9, Merlingerin, Germany). Triplicate groups of 800 nos. of 2-day post hatchling (2-DPH; mean weight 0.003±0.0001 g; mean total length 0.51±0.02 cm). The larvae were stocked into 25-L rearing aquariums for photoperiod study.

### Larval growth study

All fish in the aquaria were sampled at 5-day intervals. A sub-group of 150 randomly selected fish from both experiment collectively weighted using a digital electronic balance with provision for ±0.1 mg accuracy. Besides, lengths of a sub-group of 20 randomly selected larvae from each treatment were measured. Fish that disappeared were supposed to be victims of cannibalism. At the end of the experiment, fishes from each tank were weighed (mg) and counted. Growth parameters were calculated based on the following formulae:

Length increment (cm) = Final length – initial length

Body Weight gain (BWG) % =100 [(final weight – initial weight)/ initial weight]

Specific growth rate (SGR) % = 100[{ln (final weight) – ln (initial weight)}/ experimental period]

Mean daily Weight gain (%) = 100 [(total final weight – total initial weight)/ days of experiment]

Fulton’s condition factor (K) = (final weight/ final length^3^)

Survival percentage (%) = 100 [Total number of harvested fish/ Total number of initial stock]

Total biomass (g) = final number of fish x mean final weight

Performance index (PI) = Survival rate (%) [(Final mean body weight – Initial mean body weight) / days of experiment]

### Survival Percentage

Survival (%) was calculated at the end of the study by counting the number of fish in each treatment and is calculated as follows:

Survival (%) = Number of surviving fish / Total number of larvae stocked x 100

### Statistical analysis

The experimental data were statistically analyzed through linear and quadratic orthogonal polynomial contrasts using SPSS version 16.0 (IBM, Chicago, USA) statistical software. A 5% level of probability (P < 0.05) was chosen to determine the statistically significant difference among the treatments means. Results were presented as mean ± SE (standard error).

## Acknowledgements

The authors express great thanks to the Vice Chancellor, Central Agricultural University, Imphal, India and the Dean, College of Fisheries, CAU (I), Lembucherra, Tripura, India for their support and providing the necessary research facilities.

## Competing interests

The authors declare no competing or financial interests.

## Funding

The present work is funded by the Department of Biotechnology (DBT), Ministry of Science and Technology, Government of India, under Centre of Excellence in Fisheries and Aquaculture Biotechnology (COE FAB) project.

